## Supplemental Data for "Activation of the essential kinase PDK1 by phosphoinositide-driven trans-autophosphorylation"

#### Supplementary Figure Legends

##### Supplementary Figure S1. Intact mass spectra of recombinant proteins used in this study.

(A) – (O) Intact mass spectra for constructs used in this study. Theoretical (theor.) and experimentally determined (exp.) masses are reported, as well as any post-translational modifications.

##### Supplementary Figure S2. Intact mass spectra of recombinant proteins used in this study (continued).

(A) – (D) Intact mass spectra for constructs used in this study. Theoretical (theor.) and experimentally determined (exp.) masses are reported, as well as any post-translational modifications.

##### Supplementary Figure S3. *In silico* modeling of the PDK1 kinase domain dimer.

(A) Surface conservation and surface electrostatic potential mapping onto the PDK1 kinase domain dimer model (Rosetta).

(B) Structures of MEK1 (PDB: 1s9j) and MEK2 (PDB: 1s9i) illustrating crystallographic and non-crystallographic dimers respectively. Inset: view of αG-mediated dimer interface from superposition of PDK1 kinase domain dimer model with MEK1 and MEK2 homodimers. R.m.s.d. values are over all $C_\alpha$ atoms for both chains.

(C) B-Raf and MEK1 heterodimer structure (PDB: 6pp9) with annotation of F667 (B-Raf) and F311 (MEK1) in helix αG that are equivalent to Y288 in PDK1.

(D) Comparison of AlphaFold2 predictions of the homodimeric assembly of the PDK1 kinase domain (residues Q73-T359) to the model generated by Rosetta SymmDock. AlphaFold2 was run with and without template matching, and the ten resulting models (five from each run) superimposed on the reference Rosetta model over all Cα atoms.

#### Supplementary Figure S4. *In silico* modeling of the PDK1 kinase domain dimer.

(A) Sequence alignment of the αG helix in PDK1 orthologs spanning more than 900My of evolution.

(B) Sequence alignment of the αG helix in 26 reported or putative PDK1 substrates (all human sequences).

(C) Crystal structure of PDK1 Y288G (residues 51-359) (PDB : 3hrc). Inset: superposition of wild-type PDK1 (PDB: 2biy) and PDK1 Y288G structures.

#### Supplementary Figure S5. HDX-MS comparison of $PDK1^{SKD-PIF}$ with $PDK1^{SKD}$.

(A) Plot of differences in deuterium incorporation between $PDK1^{SKD}$ (reference) and $PDK1^{SKD-PIF}$ (reference). Changes in deuterium incorporation are plotted against the center of each peptide. Regions of protection in $PDK1^{SKD-PIF}$ are indicated below the plot and correspond to those mapped in Figure 2E-H. Error bars indicate the standard deviation of three independent replicates. Red data points indicate increases or decreases in exchange that passed the three significance criteria.

#### Supplementary Figure S6. A hydrophobic motif in PDK1 drives trans-autophosphorylation.

(A) Alignment of the kinase-PH interdomain linker of PDK1 isoforms from distantly related species. The conserved kinase domain extension and the N-bud part of the PH domain are highlighted with red boxes.

(B) Alignment of the hydrophobic motif sequence of human PDK1 identified in this study with the hydrophobic motifs of 58 other AGC kinases. Conservation and consensus sequences are indicated below the alignment.

#### Supplementary Figure S7. PDK1 is autoinhibited by its PH domain.

(A) Plot of differences in deuterium incorporation between $PDK1^{FL}$ (reference) and $PDK1^{SKD}$. Changes in deuterium incorporation are plotted against the center of each peptide. Regions of exposure in $PDK1^{SKD\text{-}PIF}$ are indicated above the plot and correspond to those mapped in Figure 4B. Error bars indicate the standard deviation of three independent replicates. Red data points indicate increases or decreases in exchange that passed the three significance criteria.

(B) Plot of differences in deuterium incorporation between $PDK1^{FL}$ (reference) and $PDK1^{PH}$. Changes in deuterium incorporation are plotted against the center of each peptide. Regions of exposure in $PDK1^{PH}$ are indicated above the plot and correspond to those mapped in Figure 4C. Error bars indicate the standard deviation of three independent replicates. Red data points indicate increases or decreases in exchange that passed the three significance criteria.

(C) PDK1 PH domain surface conservation mapped on the PH domain structure (PDB: 1w1d).

(D) PDK1 PH domain electrostatic surface potential mapped on the PH domain structure (PDB: 1w1d).

(E) PDK1 PH domain structure (PDB: 1w1d) with the N-bud highlighted in orange and annotated T513 and K465 residues.

#### Supplementary Figure S8. PDK1 is autoinhibited by its PH domain.

(A) – (C) Relative deuterium incorporation over time for peptides covering the $PIP_3$ binding site.

(D) – (F) Relative deuterium incorporation over time for peptides covering the conserved surface that includes T513.

### Figure S1. Intact mass spectra of recombinant proteins used in this study.

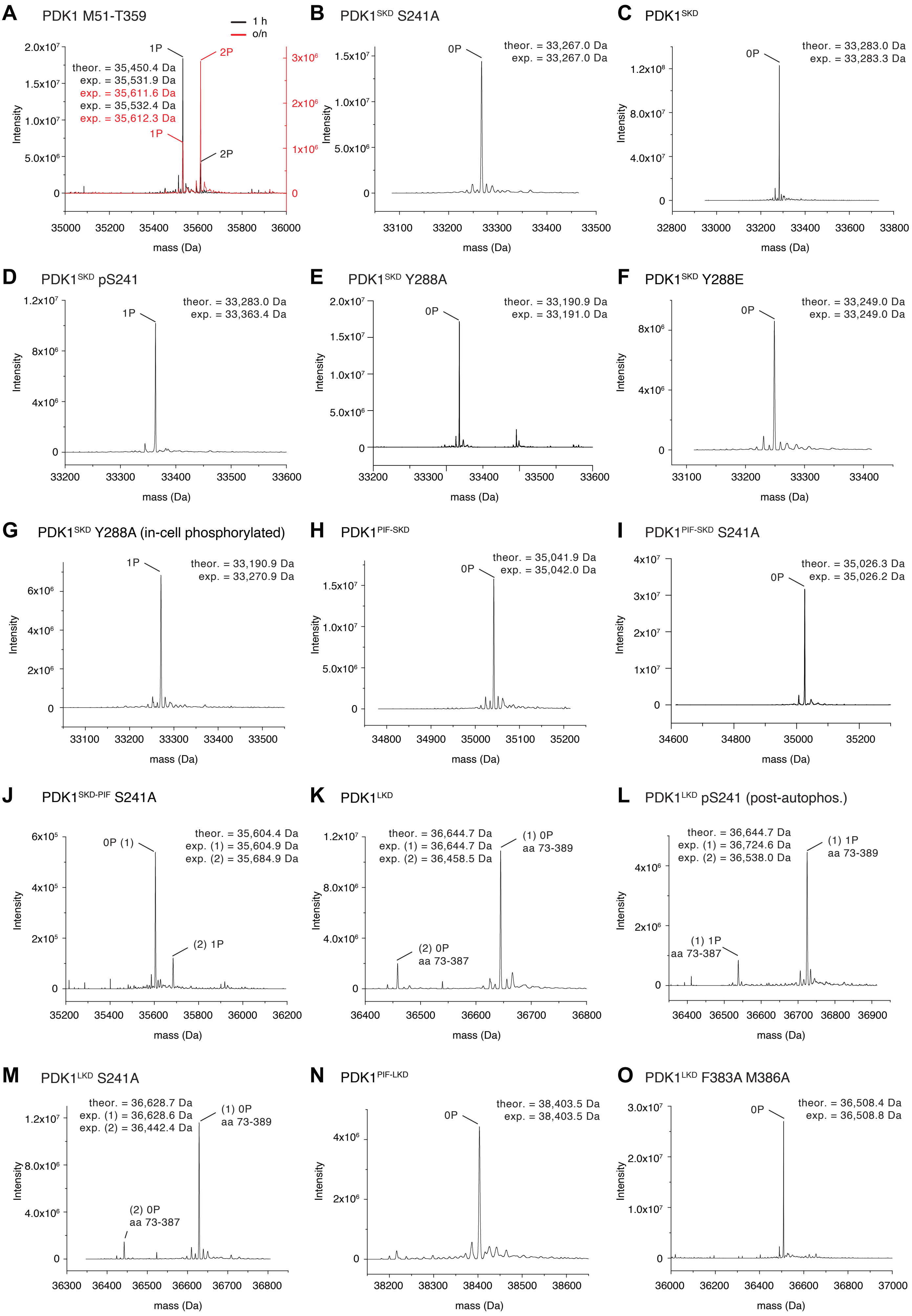

### Figure S2. Intact mass spectra of recombinant proteins used in this study (continued).

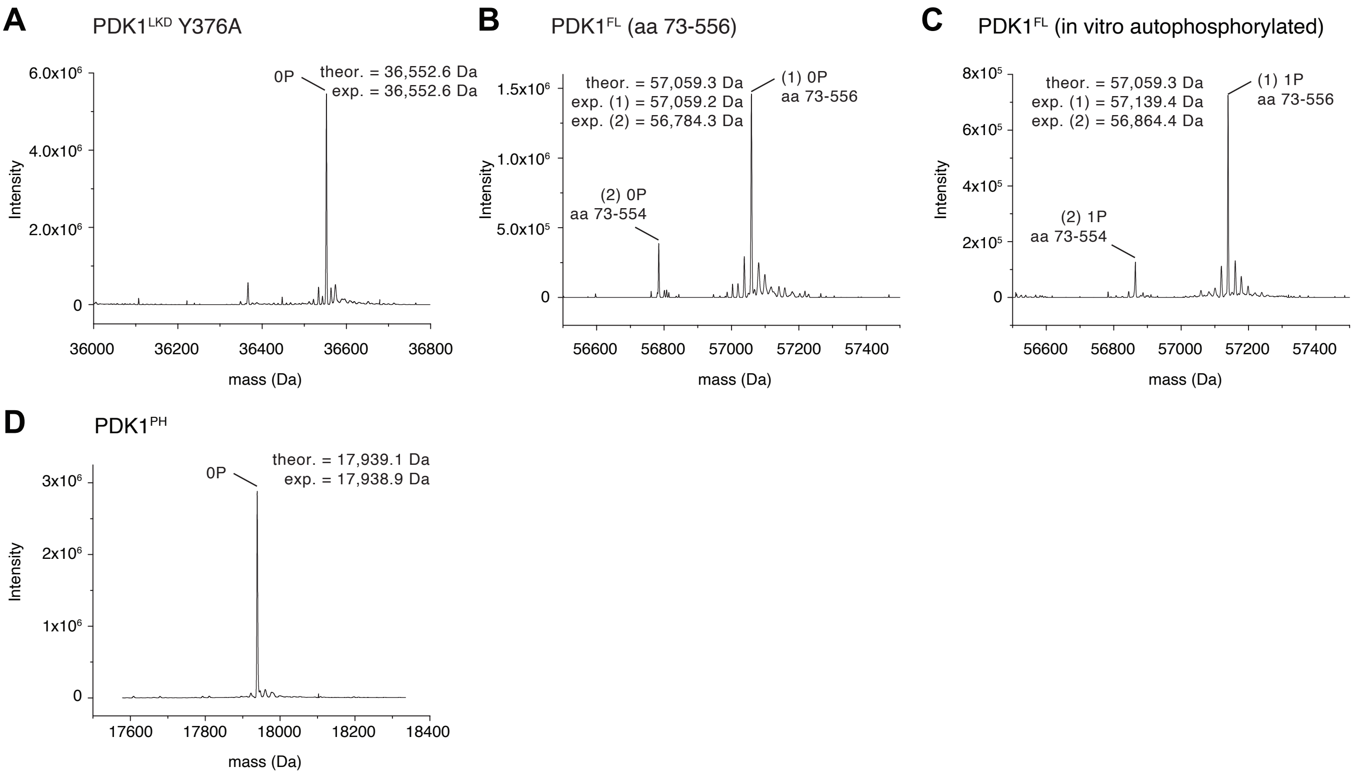

**Figure S3. *In silico* modeling of the PDK1 kinase domain dimer.**

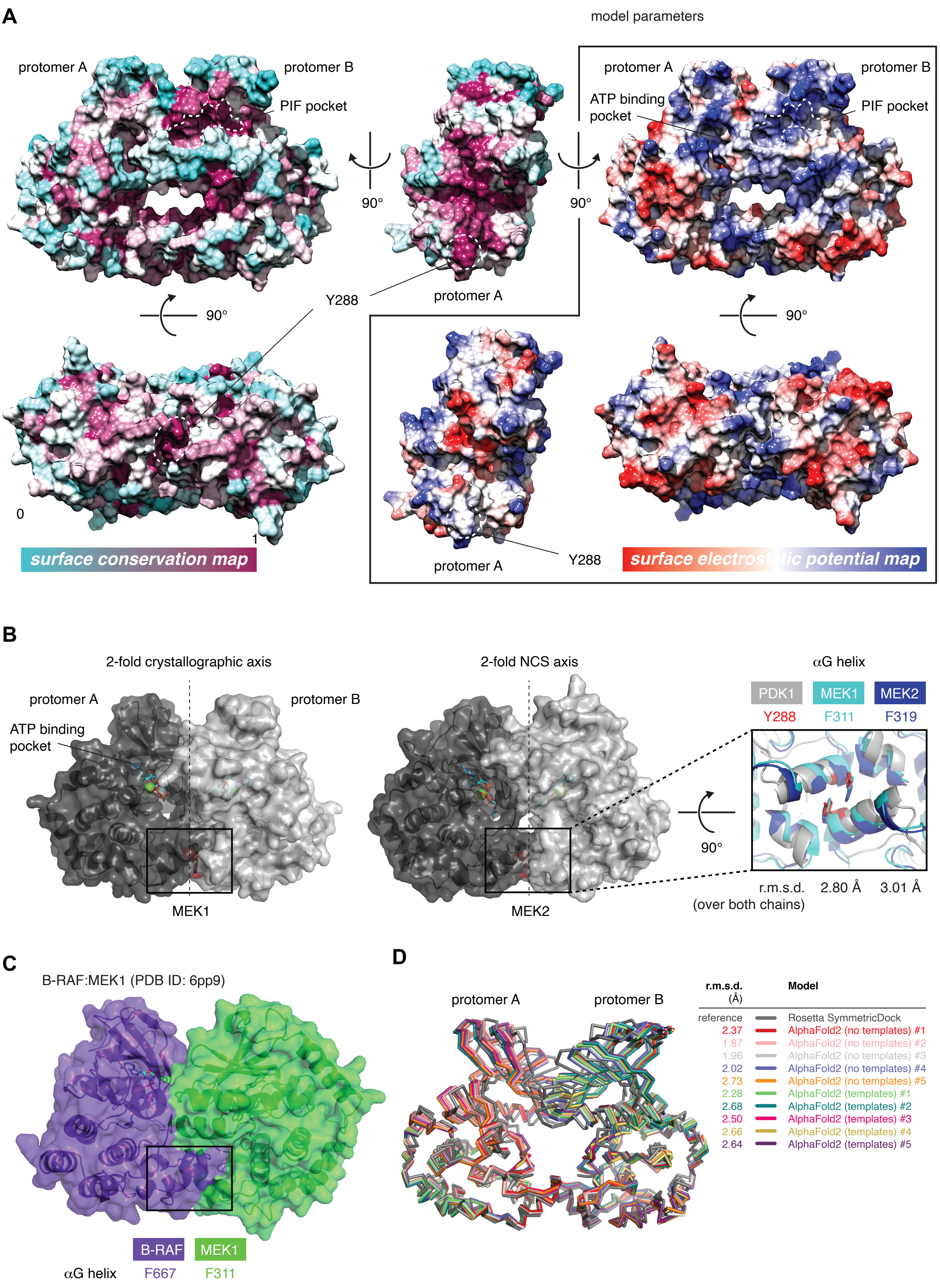

**Figure S4.** ***In silico*** **modeling of the PDK1 kinase domain dimer.**

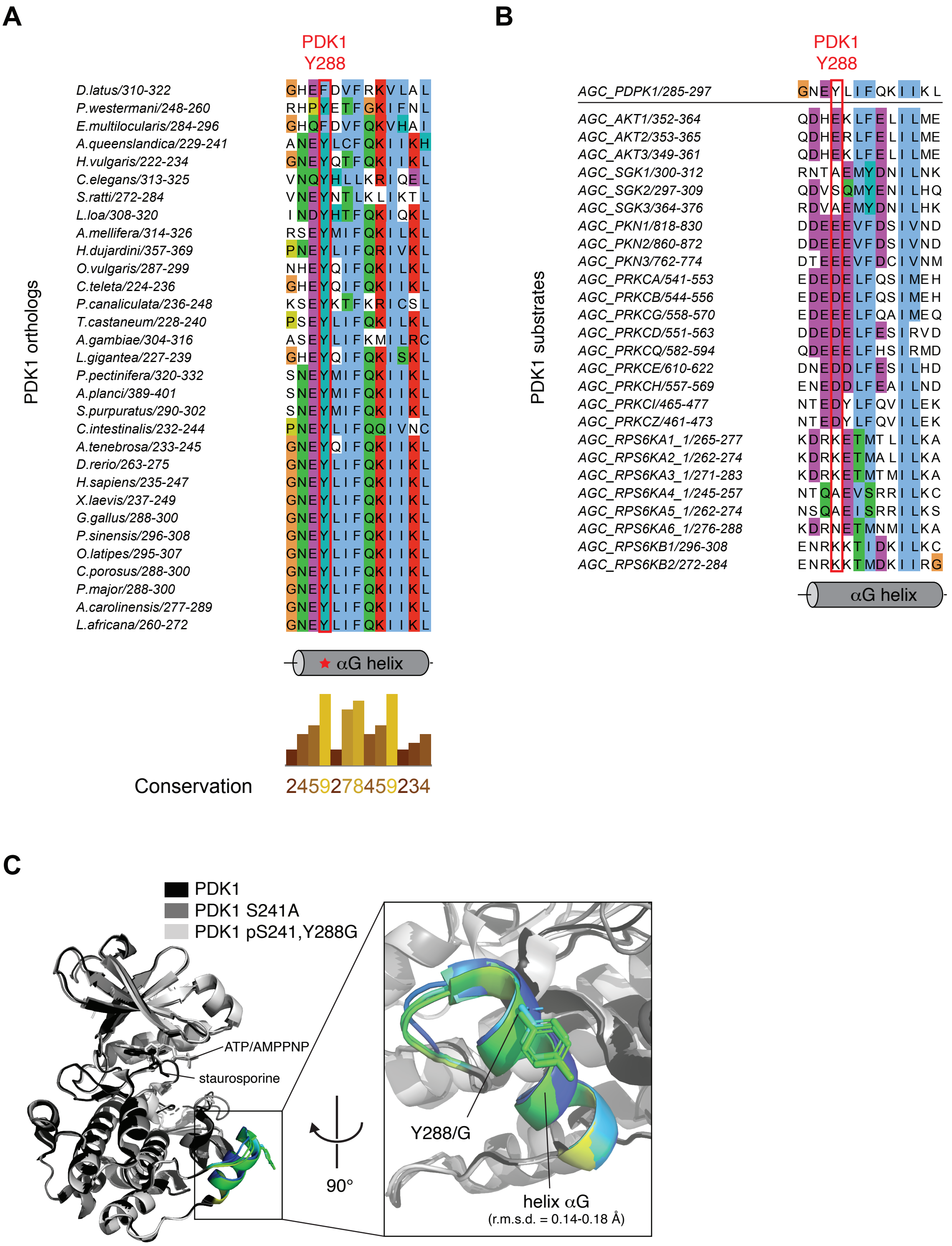

**Figure S5. HDX-MS comparison of PDK1$^{SKD\text{-}PIF}$ with PDK1$^{SKD}$.**

**A**

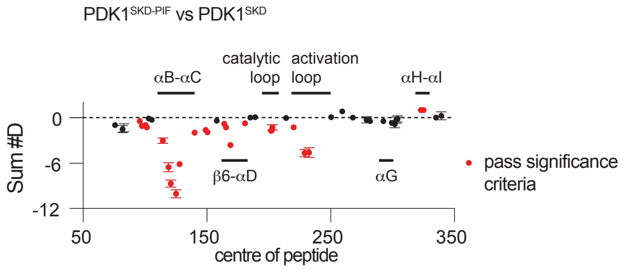

### Figure S6. A hydrophobic motif in PDK1 drives *trans*-autophosphorylation.

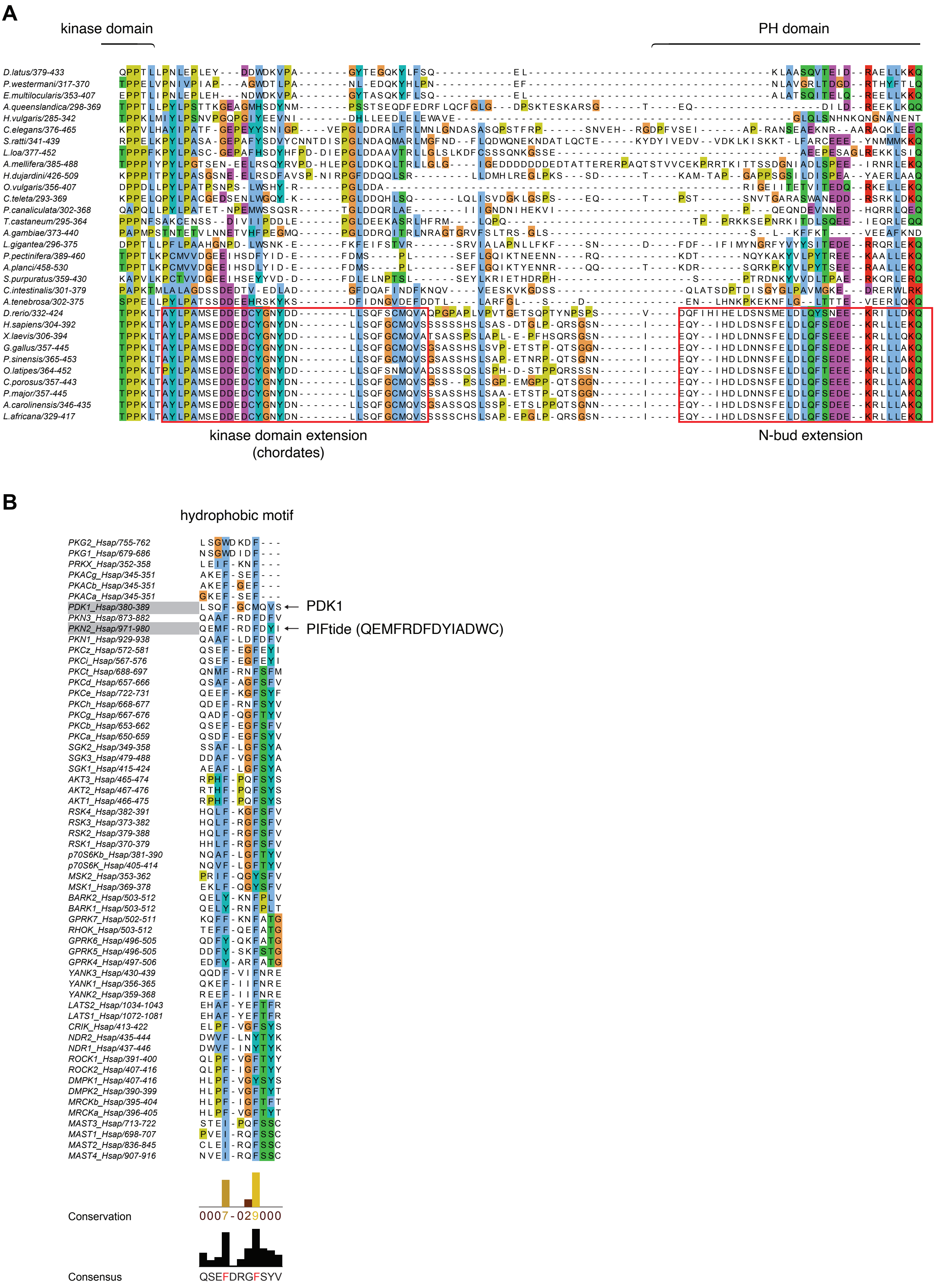

#### Figure S7. PDK1 is autoinhibited by its PH domain.

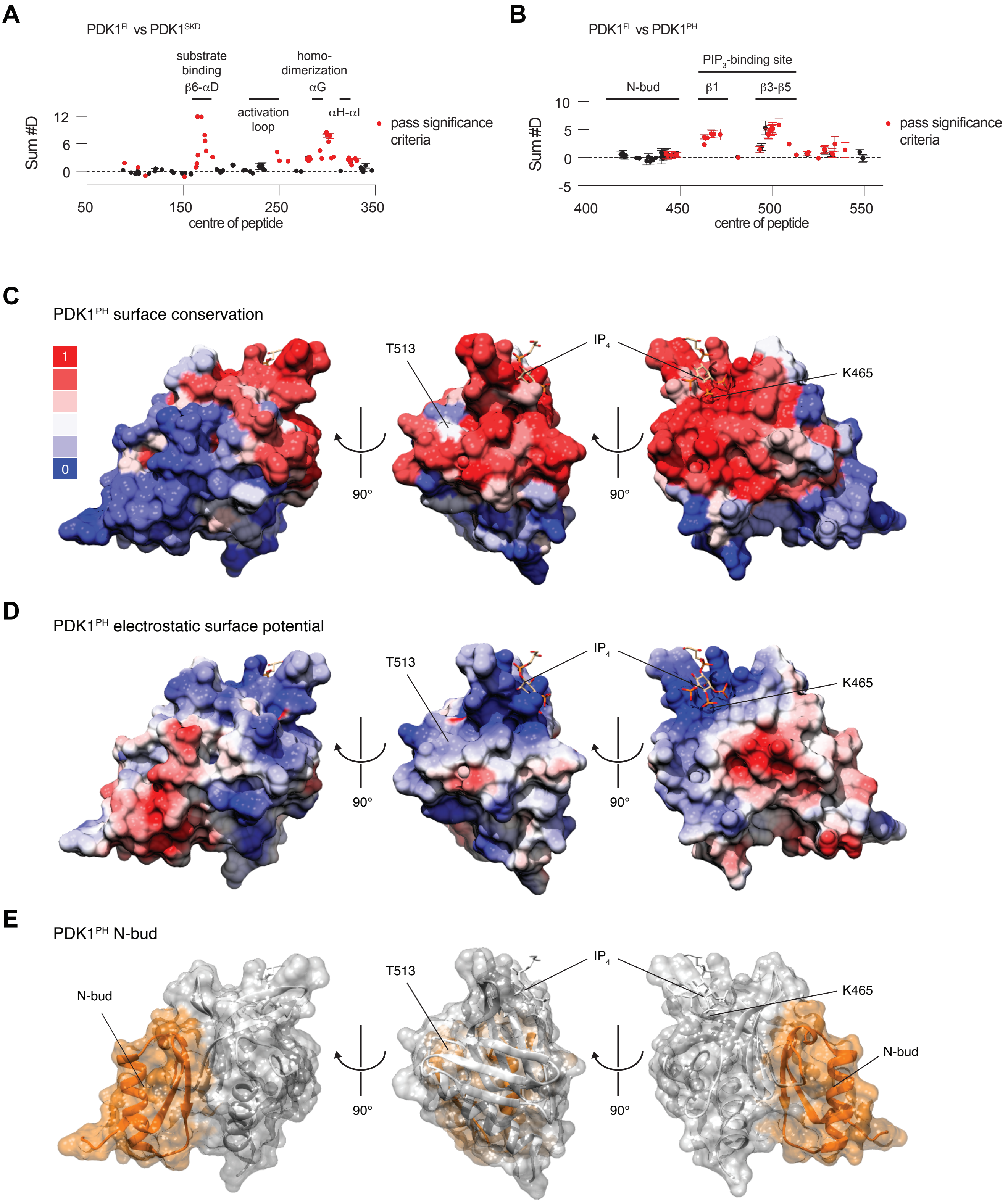

### Figure S8. PDK1 is autoinhibited by its PH domain.

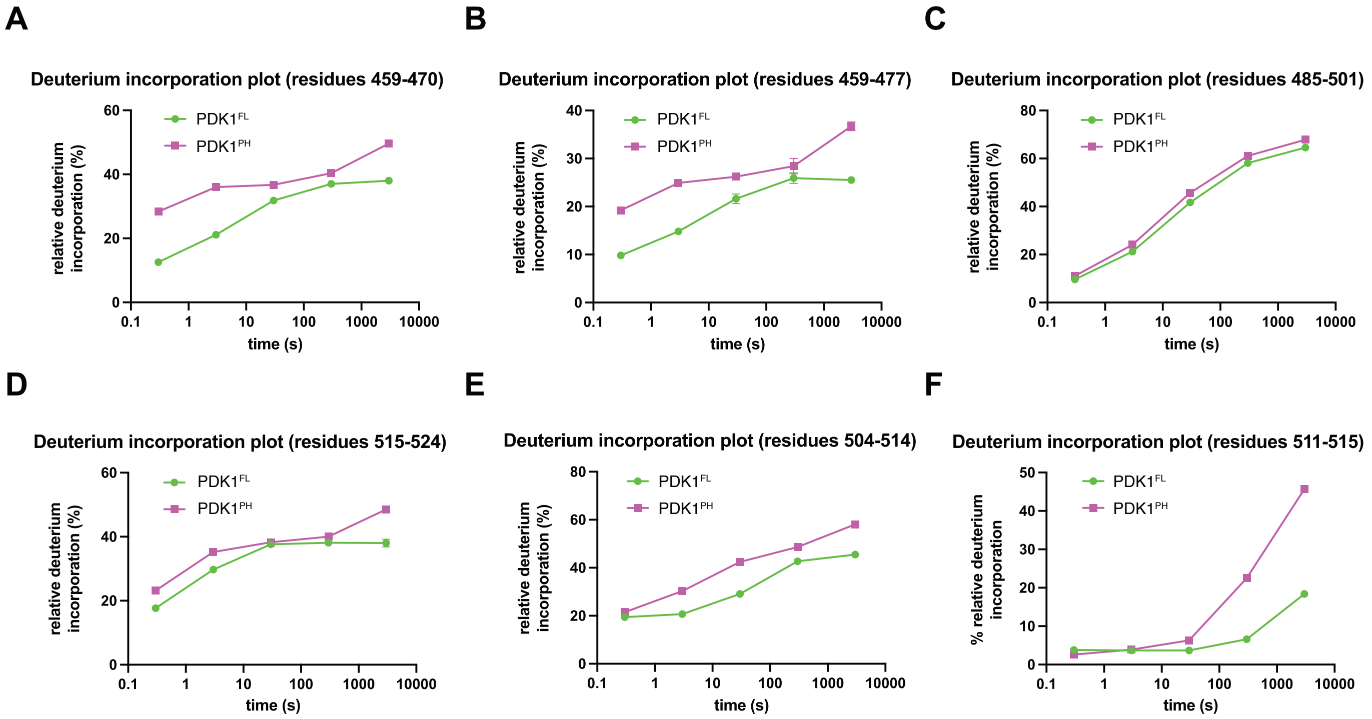

Supplementary Table 1. Constructs employed in this study.

| Construct base | Nomenclature | Description of construct |
|---|---|---|
| PDK1 kinase domain (73-359) | $PDK1^{SKD}$ | Wild-type short kinase domain |
| | $PDK1^{SKD}$ S241A | Non-phosphorylatable variant |
| | $PDK1^{SKD}$ Y288A | Dimer-disrupting mutant |
| | $PDK1^{SKD}$ Y288E | Dimer-disrupting mutant |
| | $PDK1^{PIF\text{-}SKD}$ | N-terminal PIFtide fusion (SKD) |
| | $PDK1^{SKD\text{-}PIF}$ S241A | C-terminal PIFtide fusion (SKD) |
| PDK1 long kinase domain (kinase domain + linker) (73-389) | $PDK1^{LKD}$ | Wild-type long kinase domain |
| | $PDK1^{LKD}$ S241A | Non-phosphorylatable variant |
| | $PDK1^{LKD}$ F383A/M386A | HM mutant |
| | $PDK1^{LKD}$ Y376A | 'NFD' motif mutant |
| | $PDK1^{PIF\text{-}LKD}$ | N-terminal PIFtide fusion (LKD) |
| PDK1 PH domain (408-556) | $PDK1^{PH}$ | Wild type PH domain |
| PDK1 full-length (73-556) | $PDK1^{FL}$ | Wild type full-length |
| | $PDK1^{FL}$ S241A | Non-phosphorylatable variant |

#### Supplementary Table 2. PIFtide cross-linked peptides.

| Cross-linked residues (PDK1$^{PIF-SKD}$) | Cross-linked peptide | Total peptide spectrum matches (PSM) | Dimer PSMs | Monomer PSMs |
|---|---|---|---|---|
| 1-310 | GPQPR(1)-DFDYIADW(7) | 10 | 10 | 0 |
| 1-310 | GPQPR(1)-LGSGGSSGSSGQEMFRDFDYIADW(23) | 9 | 9 | 0 |
| 103-300 | KIGSFDETCTR(1)-LGSGGSSGSSGQEMFR(13) | 1 | 1 | 0 |
| 6-310 | KKRPEDFK(1)-DFDYIADW(7) | 1 | 1 | 0 |
| 6-306 | KKRPEDFK(1)-DFDYIADW(3) | 2 | 2 | 0 |
| 1-300 | GPQPR(1)-LGSGGSSGSSGQEMFR(13) | 9 | 9 | 0 |
| 7-306 | KKRPEDFK(2)-DFDYIADW(3) | 1 | 1 | 0 |
| 50-300 | HIIKENK(4)-LGSGGSSGSSGQEMFR(13) | 1 | 1 | 0 |
| 7-310 | KKRPEDFK(2)-DFDYIADW(7) | 2 | 2 | 0 |
| | Total PSMs | 36 | 36 | 0 |

#### Supplementary Table 3. Activation loop and catalytic loop cross-linked peptides.

| Cross-linked residues (PDK1$^{PIF-SKD}$) | Cross-linked peptide | Total peptide spectrum matches (PSM) | Dimer PSMs | Monomer PSMs |
|---|---|---|---|---|
| 165-259 | VLSPESKQAR(7)-LGCEEMEGYGPLK(5) | 10 | 1 | 9 |
| 165-259 | VLSPESKQAR(7)-RLGCEEMEGYGPLK(6) | 2 | 0 | 2 |
| 165-258 | VLSPESKQAR(7)-LGCEEMEGYGPLK(4) | 3 | 0 | 3 |
| 165-186 | VLSPESKQAR(7)-ANAFVGTAQYVSPELLTEK(18) | 12 | 0 | 12 |
| 20-165 | ILGEGSFSTVVLAR(4)-VLSPESKQAR(7) | 5 | 0 | 5 |
| 165-217 | VLSPESKQAR(7)-AGNEYLIFQK(4) | 9 | 1 | 8 |
| 53-163 | ENKVPYVTR(3)-VLSPESK(5) | 3 | 0 | 3 |
| 165-182 | VLSPESKQAR(7)-ANAFVGTAQYVSPELLTEK(14) | 1 | 0 | 1 |
| 163-187 | VLSPESK(5)-ANAFVGTAQYVSPELLTEKSACK(19) | 2 | 0 | 2 |
| [a]135-137 | DLKPENILLNEDMHIQITDFGTAK(1)(3) | 6 | 0 | 6 |
| | Total PSMs | 53 | 2 | 51 |

[a]Intra-catalytic loop cross-link.

#### Supplementary Table 4. HDX-MS data processing.

| **Data set** | **PDK1 monomer** | **PDK1 dimer** | **PDK1 FL** | **PDK1 SKD** | **PDK1 FL** | **PDK1 PH** |
|---|---|---|---|---|---|---|
| HDX reaction details | %$D_2O$=66%<br>pH $_{(read)}$=7.5<br>Temp=18°C | %$D_2O$=66%<br>pH $_{(read)}$=7.5<br>Temp=18°C | %$D_2O$=89.6%<br>pH $_{(read)}$=7.5<br>Temp=18°C | %$D_2O$=89.6%<br>pH $_{(read)}$=7.5<br>Temp=18°C | %$D_2O$=81.2%<br>pH $_{(read)}$=7.5<br>Temp=18°C | %$D_2O$=81.2%<br>pH $_{(read)}$=7.5<br>Temp=18°C |
| HDX time course (seconds) | 3, 30, 300, 3000 | 3, 30, 300, 3000 | 3s at 4°C, 3, 30, 300, 3000 | 3s at 4°C, 3, 30, 300, 3000 | 3s at 4°C, 3, 30, 300, 3000 | 3s at 4°C, 3, 30, 300, 3000 |
| HDX controls | N/A | N/A | N/A | N/A | N/A | N/A |
| Back-exchange | No correction | No correction | No correction | No correction | No correction | No correction |
| Number of peptides | 46 | 46 | 102 | 102 | 69 | 69 |
| Sequence coverage | 96.9% | 96.9% | 93.1% | 93.1% | 94.8% | 94.8% |
| Average peptide length/<br>redundancy | Length=15.0<br>Redundancy=<br>2.4 | Length=15.0<br>Redundancy=<br>2.4 | Length=13.0<br>Redundancy=<br>4.6 | Length=13.0<br>Redundancy=<br>4.6 | Length=18.4<br>Redundancy=<br>8.3 | Length=18.4<br>Redundancy=<br>8.3 |
| Replicates | 3 | 3 | 3 | 3 | 3 (2 for 3000s) | 3 (2 for 30s) |
| Repeatability | Average StDev=0.6% | Average StDev=0.5% | Average StDev=0.6% | Average StDev=0.5% | Average StDev=0.6% | Average StDev=0.7% |
| Significant differences in HDX | ≥5% and ≥0.4 Da and unpaired t-test ≤0.01 | ≥5% and ≥0.4 Da and unpaired t-test ≤0.01 | ≥5% and ≥0.4 Da and unpaired t-test ≤0.01 | ≥5% and ≥0.4 Da and unpaired t-test ≤0.01 | ≥5% and ≥0.4 Da and unpaired t-test ≤0.01 | ≥5% and ≥0.4 Da and unpaired t-test ≤0.01 |
